# Enhancement of clinical signs in C3H/HeJ mice vaccinated with a highly immunogenic *Leptospira* methyl-accepting chemotaxis protein following challenge

**DOI:** 10.1101/2024.04.17.590016

**Authors:** Liana Nunes Barbosa, Alejandro LIanes, Swetha Madesh, Bryanna Nicole Fayne, Kalvis Brangulis, Sarah C. Linn-Peirano, Sreekumari Rajeev

**Affiliations:** Department of Biomedical and Diagnostic Sciences, College of Veterinary Medicine, University of Tennessee, Knoxville, Tennessee, United States; Centro de Biología Celular y Molecular de Enfermedades, Instituto de Investigaciones Científicas y Servicios de Alta Tecnología (INDICASAT AIP), Panama City, Panama; Latvian Biomedical Research and Study Centre, Riga, Latvia

**Author notes:** Corresponding author (SR).

## Abstract

Leptospirosis is the most widespread zoonosis and a life-threating disease of humans and animals. Licensed killed whole-cell vaccines are available for animals; however, they do not offer heterologous protection, do not induce a long-term protection, or prevent renal colonization. In this study, we characterized an immunogenic *Leptospira* methyl-accepting chemotaxis protein (MCP) identified through a reverse vaccinology approach, predicted its structure, and tested the protective efficacy of a recombinant MCP fragment in the C3H/HeJ mice model. The predicted structure of the full-length MCP revealed an architecture typical for topology class I MCPs. A single dose of MCP vaccine elicited a significant IgG antibody response in immunized mice compared to controls (*P* < 0.0001), especially the IgG1 and IgG2a subclasses. The vaccination with MCP despite eliciting a robust immune response, did not protect mice from disease and renal colonization. However, survival curves were significantly different between groups, and the MCP vaccinated group developed clinical signs faster than the control group. There were differences in gross and histopathological changes between the MCP vaccinated and control groups. The factors leading to enhanced disease process in vaccinated animals needs further investigation. We speculate that anti-MCP antibodies may block the MCP signaling cascade and may limit chemotaxis, preventing *Leptospira* from reaching its destination, but facilitating its maintenance and replication in the blood stream. Such a phenomenon may exist in endemic areas where humans are highly exposed to *Leptospira* antigens, and the presence of antibodies might lead to disease enhancement. The role of this protein in *Leptospira* pathogenesis should be further evaluated to comprehend the lack of protection and potential exacerbation of the disease process. The absence of immune correlates of protection from *Leptospira* infection is still a major limitation of this field and efforts to gather this knowledge is needed.

**Author summary:** Leptospirosis is one of the underrecognized and neglected diseases of humans and animals. The presence of numerous *Leptospira* species/serovars infecting a broad range of animal reservoirs, and the resulting environmental contamination, makes control and prevention a cumbersome task. The bacterin-based vaccines available for animals do not offer protection against disease or renal colonization. A broader cross-protective vaccine is essentially needed to prevent *Leptospira* infections in humans and animals. Here we rationally selected a protein target based on its capacity to be recognized by antibodies of naturally infected animals and designed a recombinant vaccine. Our MCP vaccine was not effective in protecting mice from acute and chronic disease, and likely led to exacerbation of clinical signs in these animals. The development of an effective vaccine would contribute to control *Leptospira* infection in humans and animals and is important especially in low-income regions where leptospirosis is more prevalent and interventions to control the disease are not currently available.

## Introduction

Leptospirosis is a fatal disease of humans and animals and a widespread zoonosis that causes more than 1 million human infections and 60,000 deaths annually (1). The transmission is associated with direct or indirect contact with infected animals, or exposure to contaminated water or soil (2, 3). Humans and animals with clinical leptospirosis may present with febrile illness, manifest severe forms of disease as the Weil’s Syndrome (4) with jaundice, hemorrhage and renal failure (5, 6), or the severe pulmonary hemorrhagic syndrome (SPHS) (3, 6, 7).

Although the disease is prevalent in developing countries in humans, vaccines are not available. The current bacterin-based vaccines used in animals offer protection from disease through generating antibodies to the lipopolysaccharide unique to *Leptospira* serovars. However, these vaccines provide only short-term protection restricted to the vaccine serovars and may induce adverse reactions (3, 8, 9). The *Leptospira* genus contains 66 species described to date, with 17 species characterized as pathogenic with the ability to infect a range of mammalian species (10-12). Given the presence of numerous serovars in multiple *Leptospira* species with geographic variation in their prevalence, vaccine development is difficult.

Several *Leptospira* antigens have been tested in animal models and were able to induce protective immune responses against infection in experimental studies (13, 14). However, only few targets were able to provide heterologous protection and sterile immunity (15-22). Since many of the vaccine candidates studied have not provided effective and reproducible protection against acute and chronic disease (14), new approaches to select *Leptospira* vaccine candidates are needed. Continuous improvements in bioinformatics tools provide the opportunity to identify proteins based on their potential structures, biological function, and capacity to induce protective humoral and cellular immune responses (23-25). The *in-silico* prediction of potential immunogenic *Leptospira* proteins can be used to effectively identify target molecules that can generate both humoral and/or cell mediated immune responses (26-29). In addition, studies have focused on the discovery of new vaccine candidates based on immune responses elicited by interactions of the ORFome of *L. interrogans* through microarrays (30), reverse and structural vaccinology and/or cell-surface immunoprecipitation (22, 23, 31), and the immunization with live avirulent/attenuated vaccines (32-35). These studies effectively characterized antibody profiles of patients with leptospirosis (30) and identified new potential vaccine candidates based on their interaction with sera from naturally infected hosts and vaccinated animals (22, 23, 31-35).

Recently, in a preliminary microarray-based study to evaluate the antigenicity of *in silico* predicted *L. interrogans* and *L. kirschneri* B-cell epitopes (unpublished data), we identified a repertoire of potentially immunogenic epitopes based on their reactivity to sera from dogs with clinical leptospirosis. One of the highly reactive epitopes identified was a peptide derived from a methyl-accepting chemotaxis protein (MCP) from *L. interrogans*, encoded by the gene LEP1GSC069_2151 in the genome of serovar Canicola strain Fiocruz LV133. MCPs are known to be predominant chemoreceptors in bacteria and archaea, which are involved in cell survival, pathogenesis, and biodegradation (36). MCP chemoreceptors can detect chemical changes in the environment around microorganisms, undergo reversible methylation to alter bacterial swimming behavior and to adapt to the environmental attractants and repellents (37). A typical MCP receptor consists of a ligand-binding domain, transmembrane helices, and a cytoplasmic signaling domain that interact with downstream regulatory proteins. MCPs are often localized in the poles of the cells or distributed throughout the cell body (36, 38). These proteins are further classified into four major classes (I–IV) based on their membrane topology. Membrane-embedded MCPs with periplasmic ligand-binding domains are involved in sensing extracellular signals, while cytosolic MCPs and membrane-embedded MCPs with cytoplasmic ligand-binding domains sense intracellular signals (36). Since chemotaxis is an essential process required by many pathogenic bacteria to colonize niches (36), we hypothesized that the inhibition of MCPs may have a negative effect on bacterial survival and virulence and, consequently, facilitate clearance of the bacteria from the host, limiting leptospiral dissemination and colonization of target organs. In this report, we describe the vaccination and challenge study using the MCP protein candidate in the C3H/HeJ mice model.

## Methods

### Ethical statement

All animal procedures performed in this study were approved by the University of Tennessee Institutional Animal Care and Use Committee (IACUC #2968-0523).

### Protein sequence and 3D structure prediction using AlphaFold

The full-length sequence of MCP protein coded by the gene LEP1GSC069_2151 was obtained from the GenBank database of the National Center for Biotechnology Information (NCBI) (accession number EKO69921.1). Sequence similarity among MCP proteins from *Leptospira* spp. was assessed through BLAST alignments (39) and potential orthologs from 27 representative *Leptospira* genomes were clustered using OrthoMCL (40). The prediction of signal peptides was performed by PrediSi (41), SignalP-6.0 (42) and Signal-CF software (43). AlphaFold v2.0 (44) was used to predict the 3D structure of our target MCP. Structure prediction using AlphaFold was performed using the default parameters suggested by the authors (https://github.com/deepmind/alphafold/) and was run on a computer equipped with AMD Ryzen Threadripper 2990WX 32-Core processors with 128 GB RAM and four NVIDIA GeForce RTX 2080 cards, using the full databases downloaded on 2023-10-20. For further structural analysis, only the structure predicted with the highest confidence was considered, using the predicted local-distance difference test (pLDDT) score as the confidence measure.

### Protein expression and purification

Plasmid expression vector pET-28a (+) containing the selected MCP gene fragment sequence was commercially purchased (GenScript, NJ, USA), and was inserted into competent *Escherichia coli* strain BL21 (DE3) Star cells by heat shock, following manufacturer’s instructions. The gene fragment expressed a recombinant protein of approximately 22 kDa that included the selected epitope. Recombinant *E. coli* was selected in LB medium at 37°C under agitation containing 50 µg/ml of kanamycin (Gibco™, MA, USA) as selective antibiotic. The protein expression in recombinant *E. coli* was induced by the addition of 1mM of Isopropyl ß-D-1-thiogalactopyranoside (IPTG) (Invitrogen™, MA, USA). The protein extraction and purification were performed by B-PER™ with Enzymes, Bacterial Protein Extraction Kit, Inclusion Body Solubilization Reagent and HisPur™ Ni-NTA Purification Kit, following manufacturer’s instructions (ThermoFisher Scientific, MA, USA). The purified protein was quantified by Qubit™ 4 fluorometer (Invitrogen™, MA, USA). The molecular masses of recombinant protein were evaluated by 1-D sodium dodecyl sulphate-polyacrylamide gel electrophoresis (SDS-PAGE) and western blot (45, 46). Immunoblotting was performed using Thermo Scientific™ Spectra™ Multicolor Broad Range Protein Ladder (ThermoFisher, MA, USA), and *E. coli* and *L. interrogans* bacterial extracts as controls. The recombinant protein was transferred to 0.45 μm nitrocellulose membranes using the Trans-Blot® Semi-Dry Transfer Cell (Bio-Rad, CA, USA), and bands were confirmed by 0.1% Amido black staining solution (ThermoFisher Scientific, MA, USA). The membranes were blocked in EveryBlot blocking buffer (Bio-Rad, CA, USA) for 1 h at room temperature, followed by incubation for 1 h at room temperature with primary antibodies purified from dogs naturally infected by *Leptospira,* diluted to 1:200 in EveryBlot blocking buffer. A second 1 h incubation at room temperature was performed with secondary anti-Histidine-Tag antibody (ABclonal, MA, USA) and/or anti-Dog IgG antibody (Sigma-Aldrich, MO, USA) diluted to 1:5000 in EveryBlot blocking buffer. The reactions were observed using the Thermo Scientific™ SuperSignal™ West Pico PLUS Chemiluminescent Substrate (ThermoFisher Scientific, MA, USA) in the iBright™ CL750 Imaging System (ThermoFisher Scientific, MA, USA). The purified protein was stored at -80°C until further use.

### Vaccination and challenge

3-4-weeks-old male and female C3H/HeJ mice (Jackson Laboratory, ME, USA) were housed in groups of 4 in polysulfone cages (29 cm × 19 cm × 12 cm) with soft corncob bedding, cotton nesting materials, wire-mesh tops, and paper-filter lids in a temperature-controlled colony room (22 ± 2°C), maintained on a 12:12 h light:dark cycle, with food and water available *ad libitum*. Mice (n = 8) were immunized three times with 50 µl of vaccine by intramuscular route at two-week interval (days 0, 14 and 28). The vaccines consisted of 15 µg of recombinant protein or sterile endotoxin-free PBS (negative control) (Adipogen Life Sciences, CA, USA) 1:1 in Alhydrogel® 2% (Invivogen, CA, USA) after gentle mixing for 10 minutes. Low passage *Leptospira interrogans* serogroup Icterohaemorrhagiae serovar Copenhageni strain Fiocruz L1-130 (47), kindly provided by Dr. Grassmann from the University of Connecticut, was grown in Difco™ *Leptospira* Medium Base EMJH supplemented with Difco™ *Leptospira* Enrichment EMJH (BD, NJ, USA) at 28-30°C. *Leptospira* were enumerated using Petroff-Hauser chamber under dark-field microscopy and cells in exponential growth were used for challenge. On the 42^nd^ day, mice were challenged by intraperitoneal route with 10^5^ of *L. interrogans*. After the challenge, urine samples were collected on alternate days, and the animals were monitored daily for 28 days (day 70). Animals were assessed daily using the scoring system approved by UT IACUC (S1 Table). The moribund animals were humanely euthanized through anesthetic overdose by isoflurane inhalation.

### Evaluation of immune response and protective efficacy against *Leptospira* infection

Blood samples collected on days 0, 14, 28, 42 and at euthanasia were used to evaluate the humoral immune responses using Indirect ELISA. Briefly, 96-well polystyrene plates (ThermoFisher Scientific, MA, USA) were coated with 25 ng/well of MCP protein diluted in 50 mM carbonate-bicarbonate coating buffer (bioworld, OH, USA), and incubated for 16 h at 4°C. After washing, the wells were blocked with 200 μl of blocking buffer containing 1x PBS (ThermoFisher Scientific, MA, USA), 0.05% of Tween 20™ (ThermoFisher Scientific, MA, USA) and 0.5% of non-fat milk (Nestlé^®^, CA, USA) at 37°C for 1 h. One hundred microliters of primary sera pools from each group diluted to 1:100 in blocking buffer was added to the wells, followed by incubation for 1 h at 37°C. The plates were incubated with secondary antibodies peroxidase conjugated anti-Mice IgG, IgG1, IgG2a and IgG3 (Southern Biotech™, AL, USA) diluted to 1:5000 in blocking buffer for 1 h at 37°C. The plates were washed three times with 1x PBS plus 0.05% of Tween 20™ between the incubation steps. Antigen-antibody reactions were developed by the addition of 1-Step™ TMB ELISA Substrate Solution (Invitrogen™, MA, USA) followed by ELISA Stop Solution (ThermoFisher Scientific, MA, USA). The plates were read using BioTek 800TS absorbance reader at 450 nm (Agilent, CA, USA). Antibodies titers against MCP protein were also determined by serial dilutions of pooled sera.

The vaccine efficacies were calculated based on the number of animal survivors in the vaccinated group compared to the negative control group (48). Colonization of kidneys, lungs, livers, spleens and hearts were assessed by bacterial culture (49) and/or Quantitative real-time PCR (qPCR). The DNA was extracted from urine, kidney, liver, spleen and heart tissues using the Quick-DNA Miniprep Plus Kit (Zymo Research, CA, USA), and a qPCR targeting the *Leptospira lipl32* gene was performed as previously described (50). The formalin fixed tissues were processed for histopathological assessment with routine hematoxylin and eosin (H&E) staining and evaluated microscopically by two American College of Veterinary Pathologists (ACVP) board-certified veterinary pathologists (SR, SLP).

### Statistical analysis

We used GraphPad Prism v.10 (San Diego, CA, USA) for all the statistical analyses. Survival data were plotted using the Kaplan–Meier method, and comparisons between treatment groups were made using the Log-rank Mantel-Cox and Gehan-Breslow-Wilcoxon tests. Vaccine efficacy between groups were calculated by the Fisher’s exact test (*two-tailed*). Antibody levels were compared using *two-way* ANOVA (*Turkey’s multiple comparisons* and *Dunnett’s test*). A *P-value* of ≤ 0.05 was set to assess the statistical significance for all the analyses performed.

## Results

### Expression and characterization of the MCP protein

The MCP protein used in this study is encoded by the gene LEP1GSC069_2151 in the genome of *L. interrogans* serovar Canicola strain Fiocruz LV133. This protein has 846 amino acids with an expected molecular weight of 95 kDa and was not predicted to contain a signal peptide, suggesting it is not a secreted protein. The overall sequence identity between 27 potential orthologs genes from *Leptospira* (P1, P2 and S1 clades) ranged from 31% to 100%, suggesting a high degree of conservation within P1 clade. Potential divergent orthologs also appear to be present in P2 clade (*L. licerasiae*), and the saprophytic species *L. biflexa* from S1 clade (S2 Table) (10). The structure of our target MCP, predicted from its full-length sequence with AlphaFold, revealed an architecture typical of topology class I MCPs (Fig 1A). MCPs are known to be organized as homodimers where the coiled-coil α-helices from the cytosolic signaling domains form a supercoiled four-helical bundle (51). Topology class I MCPs consists of a periplasmic ligand-binding domain, α-helical transmembrane domain, and a cytosolic HAMP domain (present in histidine kinases, adenylate cyclases, methyl-accepting proteins and phosphatases), followed by a signaling domain. Accordingly, all these domains could be identified in the predicted structure of our MCP (Fig 1B). The periplasmic ligand-binding domain (residues 217-457) is composed of a *N*-terminal α-helix that is a continuation of the α-helix (α5) of the transmembrane domain and two CACHE domains (named after the well-known functional connection of <u>ca</u>lcium channels and <u>che</u>motaxis receptors) each of which consists of an α-helix and five β-strands (Fig 1C). The cytosolic HAMP domain (residues 492-537) is a dimeric structure composed of four parallel α-helices forming a hydrophobic core region, whose structure was predicted with an average confidence score (pLDDT) of 81.6 (Fig 1D). The signaling domain, as a second cytosolic domain in MCP, is a structure approximately 230 Å long, composed of two antiparallel coiled-coil α-helices (Fig 1A). This last domain in MCPs forms a dimeric structure as both antiparallel coiled-coil α-helices from one protomer can form a supercoiled four-helical bundle with the same region in the other protomer, which could also be observed in the AlphaFold-predicted structure of our MCP (Fig 1B). The α-helices in the four-helical bundle are packed together mainly by hydrophobic interactions between the non-polar side chains while the side chains extending outward form a strongly negatively charged surface and the formed supercoiled structure showed the highest confidence score in the predicted protein structure (the average pLDDT score for residues 537-846 was 89.6) (Fig 1E and 1F).

**Fig 1.**
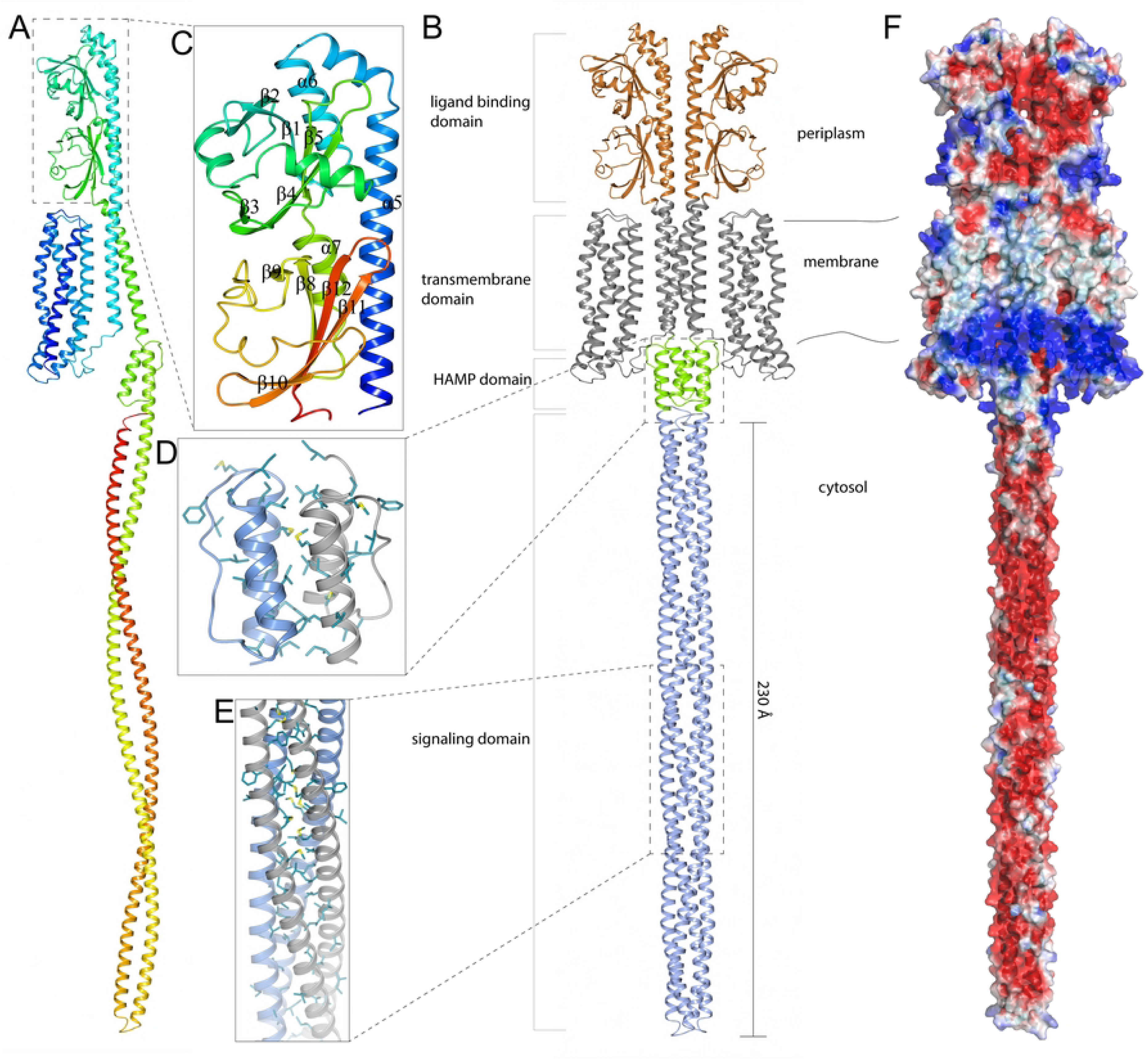
Three-dimensional structure prediction of *Leptospira* methyl-accepting chemotaxis protein LEP1GSC069_2151. (A) The predicted structure of a full-length (residues 1-846) monomeric MCP colored blue at the *N*-terminus gradually switching to red at the *C*-terminus. (B) The predicted structure of a full-length MCP homodimer. The positions of individual domains and the location of these domains in the cell are shown. (C) The periplasmic ligand-binding domain (residues 217-457) colored blue at the *N*-terminus gradually switching to red at the *C*-terminus α-helices (α5 to α7) and β-strands (β1 to β12). (D) A homodimer of the cytosolic HAMP domain (residues 492-537) where one molecule is shown in blue and the other in gray. The non-polar side chains of amino acids (Val, Leu, Ile, Phe, Met) forming the core region are illustrated as cylinders. (E) A segment of the MCP signaling domain in a homodimer structure showing the non-polar side chains that form the hydrophobic core in the supercoiled structure. One protomer is colored blue and the other gray. (F) Electrostatic surface potential of *L. interrogans* MCP. The electrostatic potentials (red = negative; blue = positive) were calculated using APBS (52). The surface contour levels were set to -1 kT/e (red) and +1 kT/e (blue).

The *C*-terminal portion of this protein, which was found to contain the seroreactive peptide epitope, was selected for further expression and evaluation as a vaccine candidate against *Leptospira* infection. The final MCP fragment contained 198 amino acids, with an expected molecular weight of 22 kDa. We purified and detected the 22 kDa MCP recombinant protein fragment in both soluble and insoluble fractions (Fig 2). The protein band was reactive to anti-Histidine-tag antibody (Fig 2A), as well as the pooled sera from infected dogs (Fig 2B), suggesting the MCP expression during *in vivo* infections.

**Fig 2.**
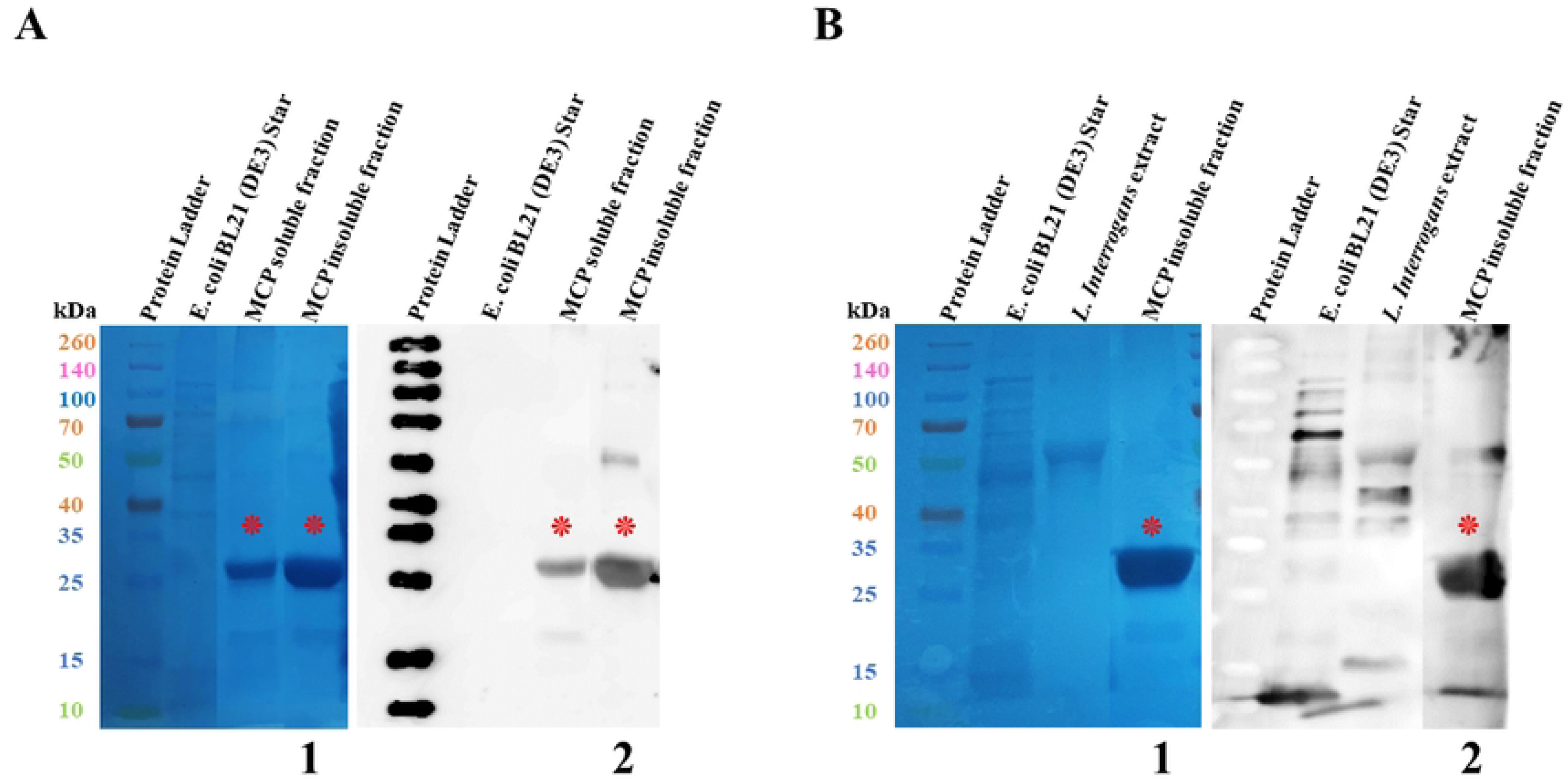
SDS-PAGE and Western blot showing the expression of soluble and insoluble forms of recombinant MCP fragment. (A) Recombinant MCP fragment with an expected molecular weight of approximately 22 kDa recognized by secondary anti-Histidine-tag antibodies. (B) Recombinant MCP fragment with an expected molecular weight of approximately 22 kDa recognized by pooled sera from dogs with clinical leptospirosis. (1) Nitrocellulose membranes after protein transfer from SDS-PAGE. (2) Chemiluminescent Western blots. (kDa) Molecular weight measured in kilodaltons. The expressed recombinant MCP fragment and antibody-antigen reactive bands are respectively indicated by red asterisks. *Leptospira interrogans* (10^8^ *Leptospira*/well) and *Escherichia coli* BL21 (DE3) Star extracts were used as controls of expression. Spectra™ Multicolor Broad Range Protein Ladder was used as reference for molecular weight.

### Immunogenicity and protection

A single dose of the MCP vaccine elicited a significant total IgG antibody response in immunized mice compared to negative controls (*P* < 0.0001) (Fig 3). Vaccination induced high levels of IgG1 and IgG2a, with minimal IgG3 immune response (Fig 4). Antibody titers to each subclass are shown in Fig 5.

**Fig 3.**
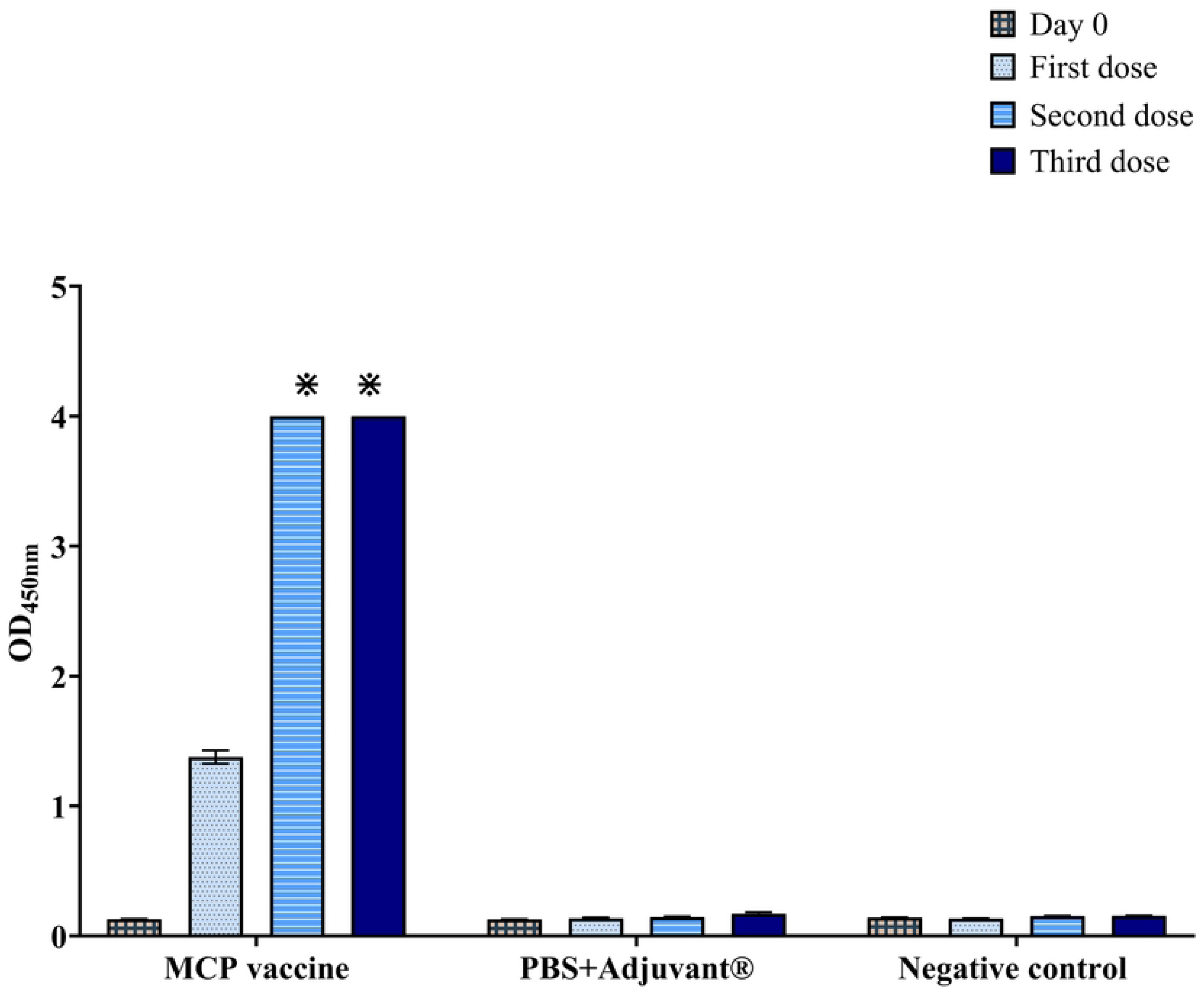
IgG antibody response elicited by immunization with MCP vaccine detected by Indirect ELISA. (OD) Optical density at 450 nanometers. (⁕) Optical density > 4.000. The columns represent the means of groups, and the bars represent the standard error. All analyses were performed in technical triplicates.

**Fig 4.**
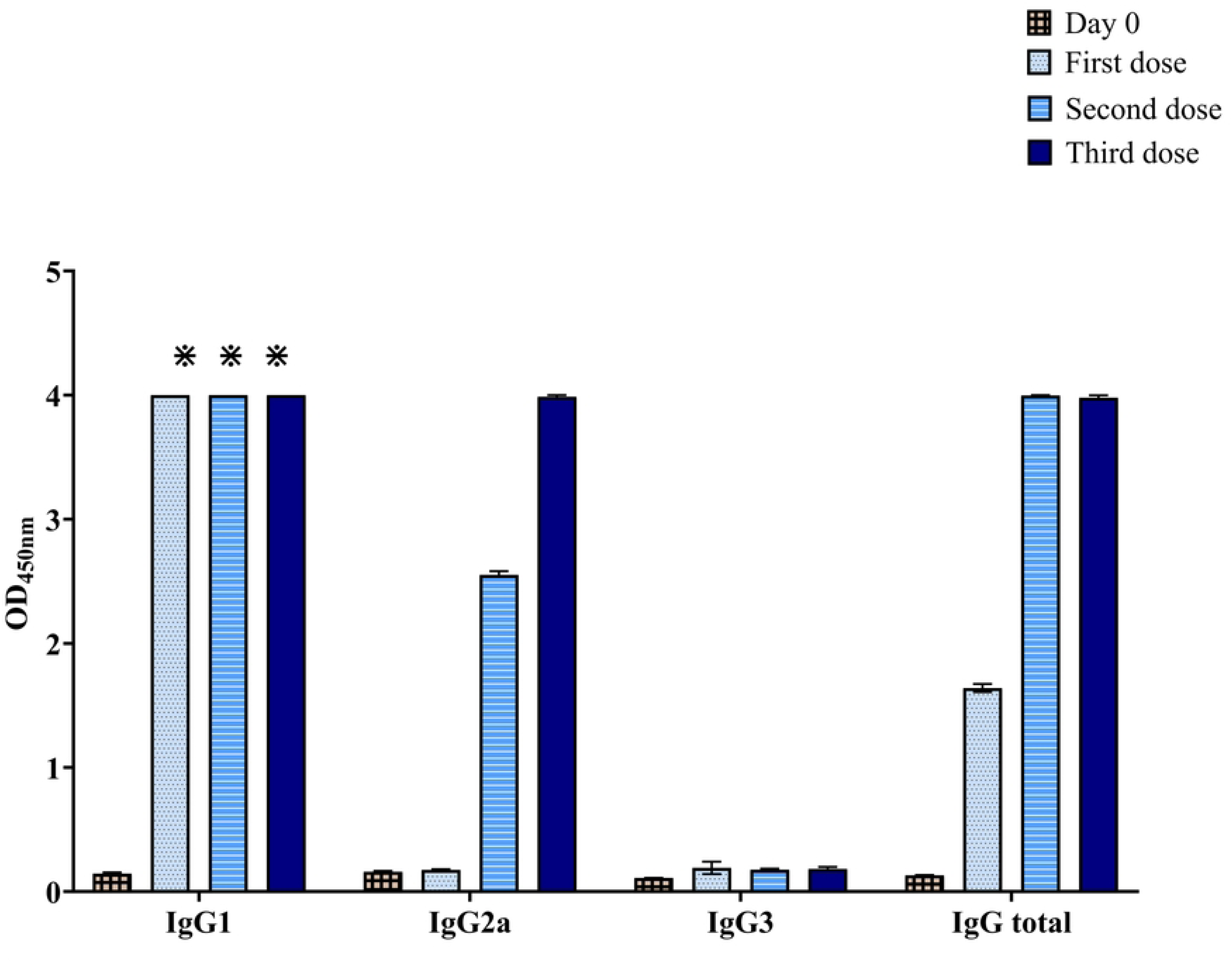
IgG subclass antibody responses induced by immunization with MCP vaccine detected by Indirect ELISA. (OD) Optical density at 450 nanometers. (⁕) Optical density > 4.000. The columns represent the means of groups, and the bars represent the standard error. All analyses were performed in technical triplicates.

**Fig 5.**
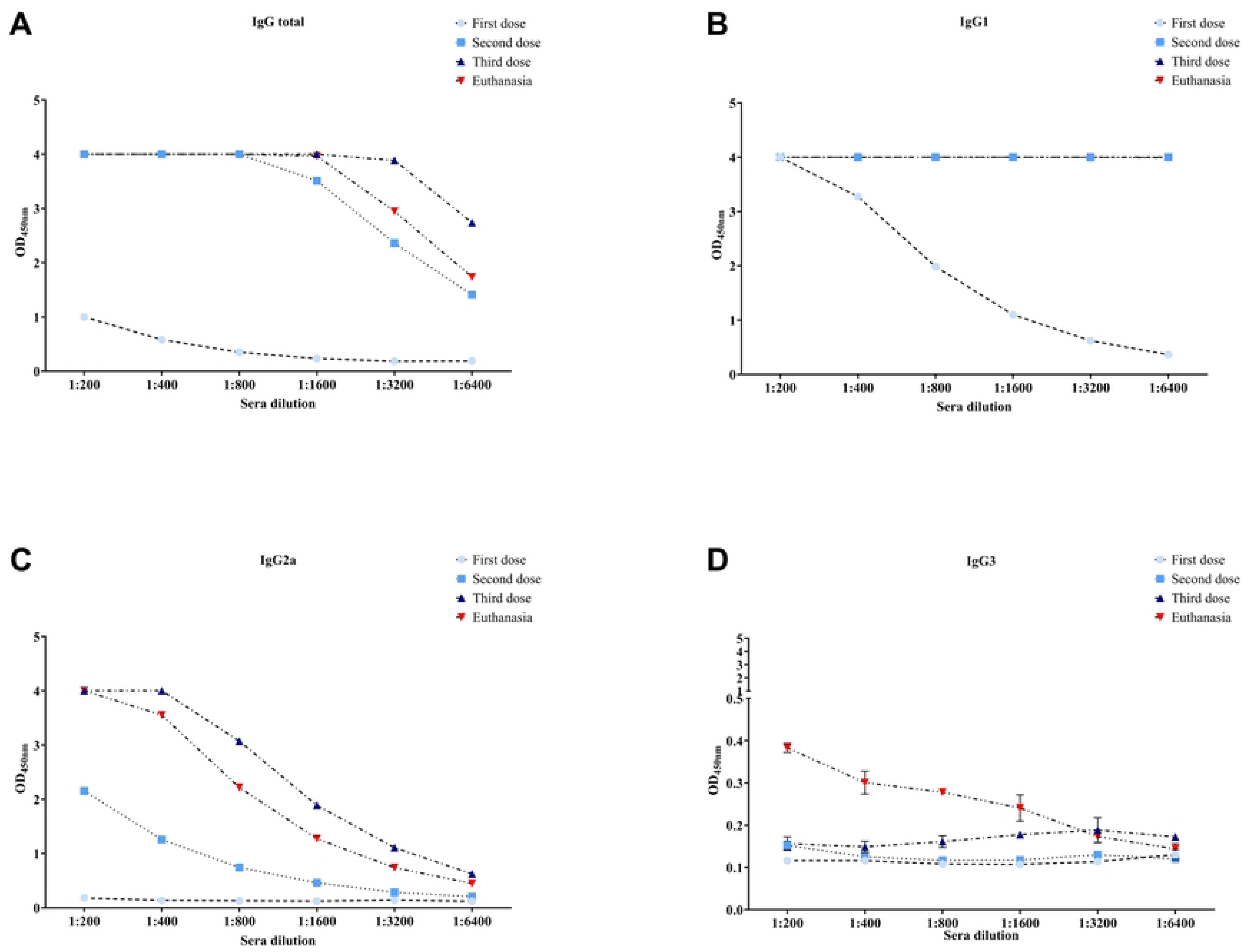
Titration of IgG subclasses induced by immunization with MCP vaccine. (OD) Optical density at 450 nanometers. (A) Total levels of IgG antibody. (B) Levels of IgG1 subclass. (C) Levels of IgG2a subclass. (D) Levels of IgG3 subclass. The bars represent the standard error. All analyses were performed in technical triplicates.

All animals in the unvaccinated/unchallenged negative control group survived and did not show any clinical signs of leptospirosis. In the MCP vaccinated group, 7/8 animals (87.5%) reached the endpoint criteria between 5-10 days, whereas 5/8 (62.5%) animals reached the endpoint between 7-10 days in the PBS+Alhydrogel® control group (Fig 6A). No significant difference in protection against acute disease was observed between MCP vaccine and the PBS+Alhydrogel® control group (*P* = 0.5692). When the survival rates curves were compared using Log-rank (Mantel-Cox) test, there was no significant difference between MCP vaccinated and control groups (*P* = 0.0570). Median days of euthanasia was lower in MCP vaccinated group (7 days) compared to control group (10 days). When Gehan-Breslow-Wilcoxen test (which gives more weight to deaths at early time points) was used, survival rates between the MCP vaccinated and control group were significantly different (*P* = 0.0361).

**Fig 6.**
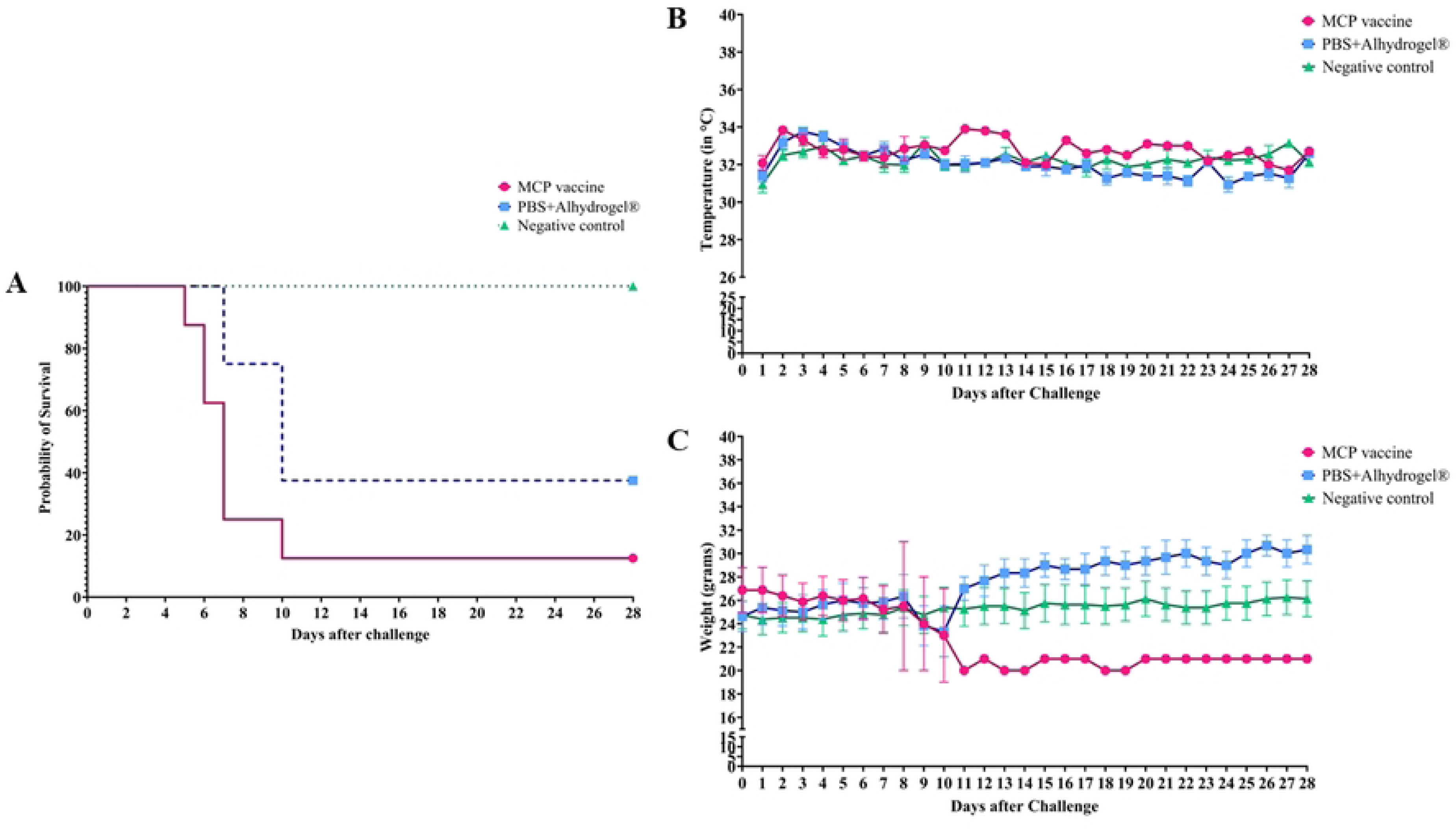
Survival rates, body weight and body surface temperatures of C3H/HeJ mice infected with pathogenic *Leptospira*. (A) Survival rates of C3H/HeJ mice from experimental groups. (B) Surface body temperatures of immunized C3H/HeJ mice after challenge with pathogenic *Leptospira* and from unvaccinated/unchallenged negative control group. (C) Weight in grams of C3H/HeJ mice from experimental groups. (°C) Celsius degrees. (g) Grams. The means of temperature and body weight were calculated only if more than one animal survived in the group. The lines represent the means of each group with standard error bars.

During the progression of the disease, typical leptospirosis-related clinical signs were observed in infected animals. The surface body temperatures of 4/8 euthanized animals in the MCP vaccinated group slowly decreased after challenge (Fig 6B), leading to hypothermia in one animal (> 4°C, day 5) and the decrease of at least 2.6°C on day 7 in the three remaining animals. In the PBS+Alhydrogel® control group, only one animal (1/8) displayed a reduction of surface body temperature of 1.6°C, which associated with other clinical signs, resulted in euthanasia on day 7 after challenge. Additionally, 4/8 euthanized animals in the MCP vaccinated group had a body weight loss of 6-18%, along with other clinical signs, resulting in the euthanasia of these animals (Fig 6C). Similarly, 3/8 animals from PBS+Alhydrogel® control group had a decrease in body weight of 18-21%. Other leptospirosis-related clinical signs observed in euthanized animals included decreased activity, hunched back, piloerection, sunken eyes, ocular discharge, and moderate to severe jaundice.

### Gross and histopathological findings

Moderate to severe jaundice was externally visible in the skin and body cavities of 5/8 animals in the MCP vaccine group, and 2/8 in the PBS+Alhydrogel® control group (Fig 7A, B and C). Splenomegaly was observed in some of the euthanized animals in both groups. In animals from both groups, the renal changes included lymphoplasmacytic interstitial nephritis, tubular degeneration, necrosis, occasionally mildly dilated tubules, and tubular hyaline casts. However, renal changes in the PBS+Alhydrogel® control group tended to be more severe and included tubular degeneration and necrosis and lymphoplasmacytic interstitial nephritis. Surviving animals in both groups, euthanized on day 28 after challenge, had chronic changes including interstitial nephritis and rare to multifocal periglomerular fibrosis. Occasionally, animals from both groups had evidence of pyelitis and neutrophilic inflammation. This is suspected to be an unrelated background lesion and may be associated with vesicoureteral reflux in this mouse strain (53). Histopathologic changes in the liver were generally unremarkable. Animals with jaundice appeared to have mild hepatocellular dissociation, and minimally decreased hepatocellular cytoplasmic swelling and clearing (decreased glycogen stores). The PBS+Alhydrogel® control group had mildly increased presence of mitotic figures. The histopathologic changes are shown in Fig 8. A summary of gross and histopathological changes observed in individual animals is shown in S3 Table.

**Fig 7.**
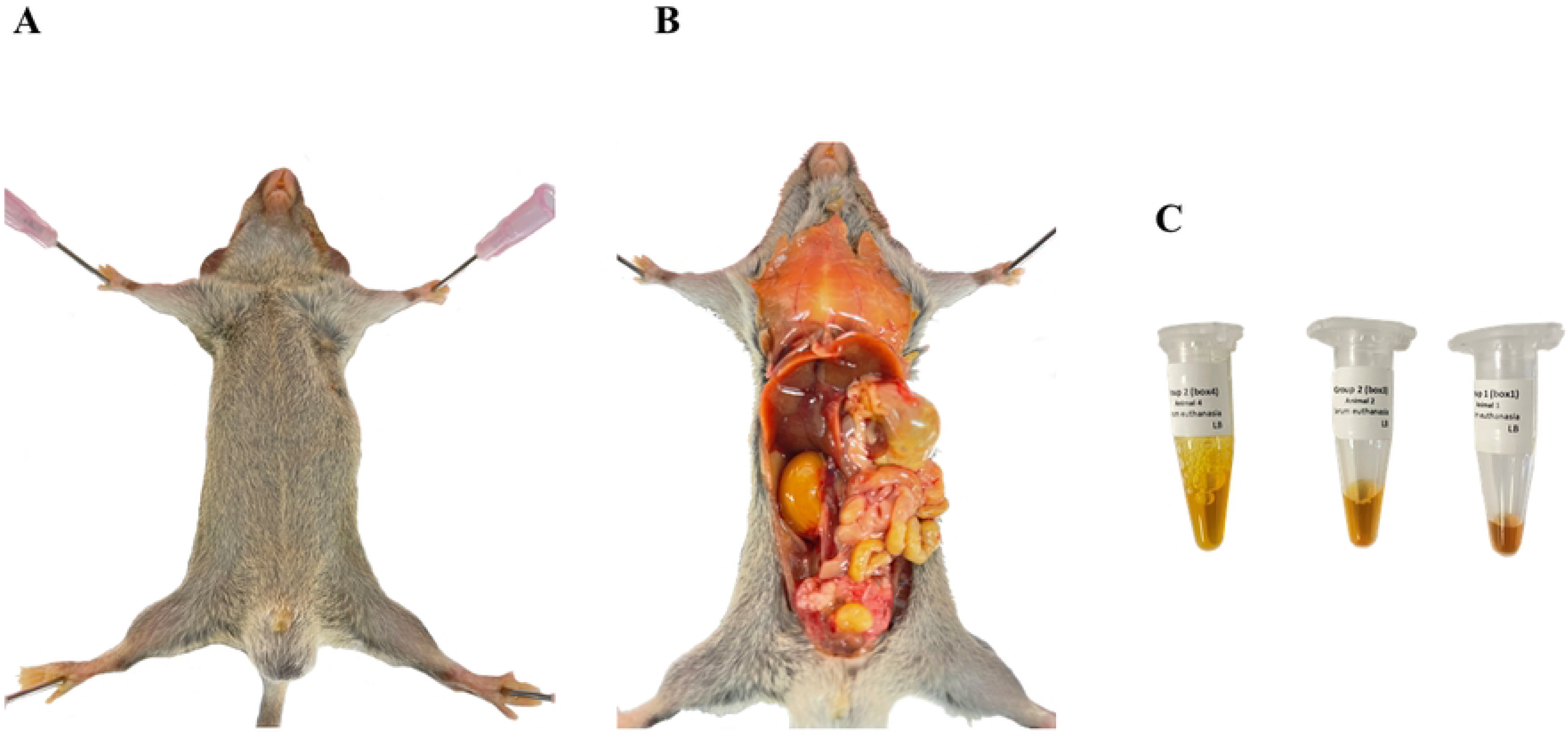
A representative image of jaundice observed in C3H/HeJ mice infected with pathogenic *Leptospira*. (A) External surface of body of infected mouse. (B) Internal organs of infected mice. (C) Sera samples collected at euthanasia of clinically ill animals.

**Fig 8.**
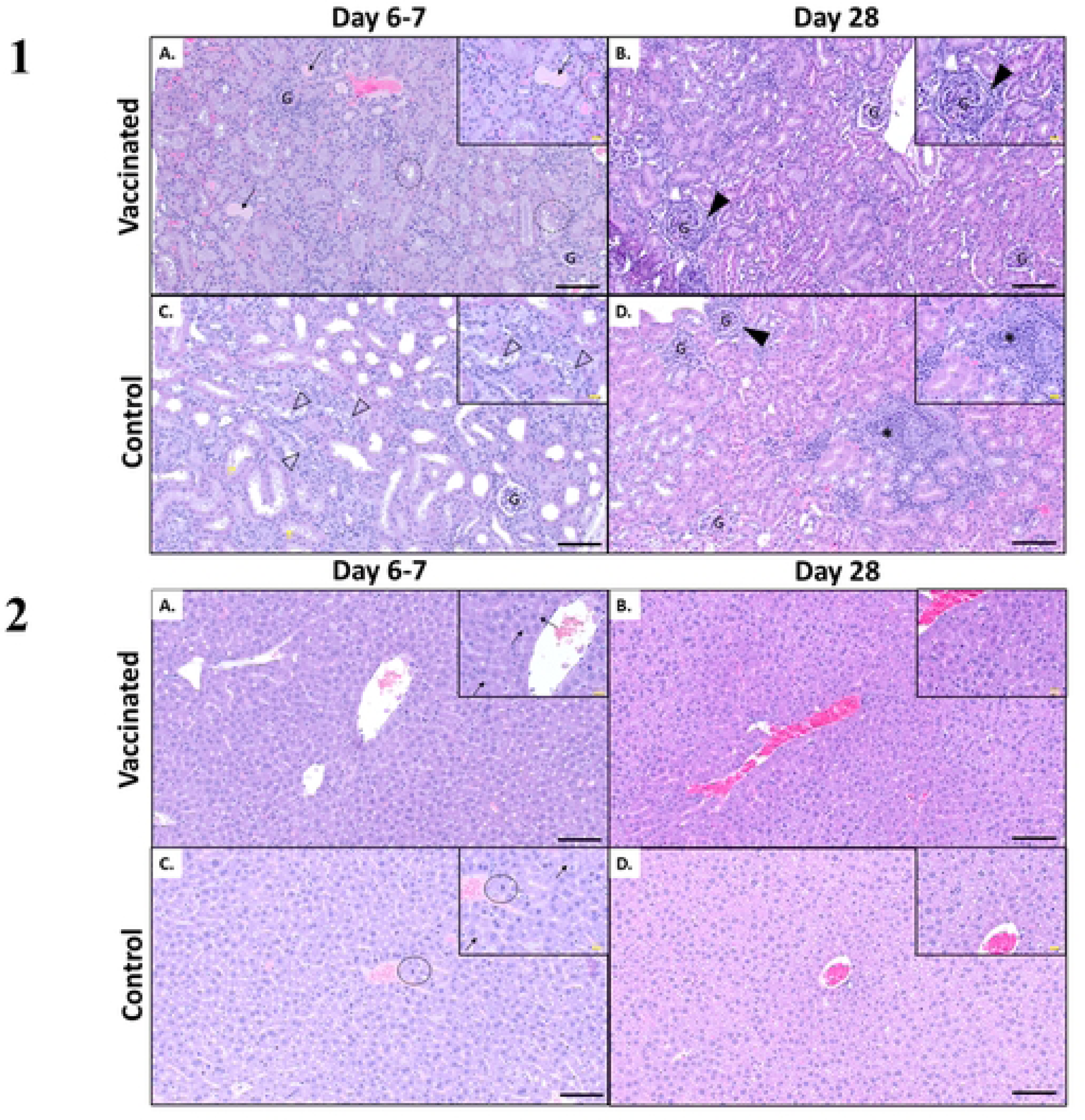
Histopathologic changes in the kidneys and livers from mice vaccinated with MCP vaccine candidate or PBS+Alhydrogel® and challenged with pathogenic *Leptospira*. (1) *Kidney.* (A and B) Representative H&E stained kidney sections from MCP vaccinated mice. (A) Euthanized on day 6 post-infection. There were minimal changes to the kidney tubules. Occasionally hyaline casts were present (black arrow). There was minimal cytoplasmic vacuolation to the proximal tubules (black dotted circles). (B) Kidneys from MCP vaccinated mouse, that survived the challenge up to 28 days post-infection, only had one region of periglomerular fibrosis (solid black arrowhead). The remaining kidney was within normal limits. (C and D) Representative H&E stained kidneys from PBS+Alhydrogel® control mice. (C) Euthanized on day 7 post-infection. There was moderate tubular ectasia (dilation) in the proximal tubules. There was acute tubular necrosis and degeneration with necrotic cells and debris within the tubular lumens (black outlined arrowhead). Tubular epithelium was attenuated (flattened) and there was cytoplasmic basophilia (degeneration). There was cytoplasmic blebbing within the proximal tubules (yellow arrow). (D) In the kidneys of mice that survived to day 28 post-infection, there were multifocal mild to moderate regions of lymphoplasmacytic interstitial inflammation (black asterisk) with loss of tubules. Occasionally, there was increased periglomerular fibrosis (solid black arrowhead). (G) Glomerulus. (2) *Liver.* (A and B) Representative H&E stained liver sections from MCP vaccinated mice. (A) Euthanized on day 6 post-infection. Hepatocytes were typically arranged in normal hepatic cord structure. Hepatocyte cytoplasm was uniformly eosinophilic with mild perinuclear clearing (black arrow) (presumptive Golgi apparatus). (B) Euthanized on day 28 post-infection. Hepatocytes were mildly disorganized from the normal hepatic cord structure. There was increased cytoplasmic swelling and clearing in the hepatocytes (glycogen). (C and D) Representative H&E stained kidneys from PBS+Alhydrogel® negative control mice. (C) Euthanized on day 7 post-infection. There was moderate hepatocellular cord dissociation. Hepatocyte cytoplasm was uniformly eosinophilic with mild perinuclear clearing (black arrow) (presumptive Golgi apparatus). There were increased mitotic figures throughout the liver (black circle). (D) Euthanized on day 28 post-infection. Hepatocytes were arranged in normal hepatic cord structure. There was increased cytoplasmic swelling and clearing in the hepatocytes (glycogen). Black scale bar = 200 µm. Yellow scale bar = 20 µm.

### *Leptospira* colonization in organs

We detected *Leptospira* DNA by qPCR in kidneys, lungs, livers, spleens, and hearts collected from challenged animals, except the heart of one animal in the PBS+Alhydrogel® group (Table 1, S3 Table). The Ct values ranged from 21.12 to 33.99, suggesting the presence of moderate to heavy *Leptospira* DNA in these samples (S3 Table). In addition, *Leptospira* DNA was also detected in urine samples taken at the euthanasia for 2/3 animals in the MCP vaccine (Ct values 27.932 and 27.805) and 5/5 animals from PBS+Alhydrogel® (Ct values from 20.15 to 28.20) groups. No *Leptospira* DNA was detected in the urine of animals from unvaccinated/unchallenged group. We recovered *Leptospira* from kidneys of 5/8 animals from the MCP immunized group and 6/8 animals from the PBS+Alhydrogel® control group. The cultures from unvaccinated/unchallenged group were negative for *Leptospira* by culture and qPCR. The summary of overall findings from this study is shown in Table 1.

**Table 1.** Summary of overall findings from the vaccination/challenge study.

| Experimental group | Protection (%) | Survivors | Days to reach endpoint (n° of animals) | Culture results | qPCR results |  |  |  |  |  |
| --- | --- | --- | --- | --- | --- | --- | --- | --- | --- | --- |
|  |  |  |  |  | Kidneys | Lungs | Spleen | Heart | Liver | Urine |
| MCP vaccine | 12.5 | 1/8 | 5-10 (7) | 5/8 | 8/8 | 8/8 | 8/8 | 8/8 | 8/8 | 2/3 |
| PBS+Alhydrogel® | 37.5 | 3/8 | 7-10 (5) | 6/8 | 8/8 | 8/8 | 8/8 | 7/8 | 8/8 | 5/5 |
| Unvaccinated/Unchallenged negative control | - | 4/4 | - | 0/4 | 0/4 | 0/4 | 0/4 | 0/4 | 0/4 | 0/4 |

(%) Percentage.
(n°) Number.
(qPCR) Quantitative real-time polymerase chain reaction.

## Discussion

In this study, we predicted the structure of a *Leptospira* methyl-accepting chemotaxis protein (MCP), identified through microarray screening, and evaluated the protective efficacy of a recombinant MCP fragment in C3H/HeJ mice. We concluded that despite the high level of humoral immune response induced, the immunization with MCP protein did not offer protection but likely worsened the clinical disease outcome.

MCPs are the most common sensing molecules found in bacteria and archaea, and they direct motility towards favorable environments through sensing chemical cues. Several classes of MCPs have been identified and the signaling process through these proteins modulates pathogen’s motile behavior and facilitate their colonization and virulence (36, 38). MCPs together with the downstream adaptor and regulatory proteins such as CheA, CheW and CheY form a chemotaxis system, the most common type of signal transduction system in bacteria to control cell movement (54-56). Three-dimensional prediction using AlphaFold revealed that the MCP fragment containing the seroreactive epitope is in the cytosolic region of the MCP protein. The reactivity of this epitope to sera from infected dogs at the early stage of infection suggests that, in fact, the cytosolic portion is somehow exposed to the immune system. The absence of reactivity of *L. interrogans* extract to these antibodies suggests low level or no expression of MCP in *in vitro* cultures. Proteins detected by antibodies from infected animals are potentially regulated during *in vivo* infections, suggesting they are important factors that might play a role during host-pathogen interactions. Further functional studies are needed to elucidate the role of MCP in *Leptospira* pathogenesis. A recent study evaluating a live-attenuated *fcpA^-^* vaccine against leptospirosis identified a total of 154 unique protein targets by immunoproteomics, including a MCP protein (35). In a *Burkholderia pseudomallei* study, a group of 6 new outer membrane proteins not reported previously as vaccine targets were identified, including a methyl-accepting chemotaxis protein III (57). These two studies suggested that MCPs are antigenic, are essential during bacterial infections and may have potential as vaccine antigens.

With the aim to develop an effective vaccine to prevent *Leptospira* infection and colonization, a number of prospective new vaccinal targets and approaches have been explored (14). Recently, there was a vaccine study performed in 13 independent experiments in hamsters using 22 new vaccine candidates composed by *Leptospira* β-barrel outer membrane proteins (22). In this study, only a few targets were able to significantly protect or increase survival of vaccinated animals. However, the reproducibility of independent experiments was a major concern (22). The authors of this paper suggest that the lack of consistency between experiments are due to the high susceptibility of the Golden Syrian hamsters to leptospirosis (9, 58, 59). The C3H/HeJ is a mice strain with a spontaneous mutation in *tlr4* gene rendering them hyporesponsive to LPS (60, 61), and has been recommended to study *Leptospira* infections (62). The lethal effect of gram-negative bacterial infections is partially related to the biological effects of lipopolysaccharides (LPS), and this model is presumed to bypass the initial acute endotoxemia and shock phase (63). However, it is also suggested that C3H/HeJ mice is somewhat immunocompromised due to this defect in the *tlr4* gene, making it highly susceptible to *Leptospira* infection (61). *Leptospira* infection in C3H/HeJ mice has been widely studied and these animals exhibit acute and chronic leptospirosis (22, 62, 64). There are also few studies that used this model to test vaccine candidates (65, 66). The infected mice exhibited many acute manifestations of leptospirosis, including jaundice, pulmonary hemorrhage, renal failure, massive splenomegaly, higher leptospiral burden in target organs and histopathological changes in lungs and kidneys (64, 67-69), which recapitulates severe leptospirosis symptoms in humans and animals (69, 70). We observed many abovementioned changes in challenged mice in our study. Nevertheless, the actual impact of *tlr4* mutation in this model is still unclear.

In our study MCP antigen elicited significant IgG antibody levels in immunized C3H/HeJ mice, predominantly IgG1 and IgG2a isotypes but minimal IgG3, despite the lack of protection. Factors associated with correlates of protection are still unknown in cases of leptospirosis (17, 19, 21, 22, 71-81). In fact, our study reinforces the need to improve our understanding of the protective immune responses against leptospirosis, especially the role of cellular immune response, which is poorly evaluated in experimental vaccines tested in the hamster model. One of the drawbacks of our study is the use of an antigen which is not surface exposed, differently than the vast literature in the last 25 years. Nonetheless, the use of *Leptospira* antigens not exposed in the outer membrane is common in the field and elsewhere. For example, a recent study evaluated recombinant fragments of a potential toxin candidate from *Leptospira* with some success (66). In another study, a leptospiral recombinase A (RecA) and a flagellar hook associated protein (FliD) induced humoral and cellular immune responses and provided a significant protection against homologous and heterologous challenge (82). Although MCP proteins are transmembrane proteins, the epitope reactivity suggests that this protein may be exposed to the immune system during natural infection and can elicit robust antibody response. However, we do not have any knowledge about the functional aspects of this protein, including its potential ligands.

In our study exacerbation of clinical signs and shorter days to euthanasia were observed in MCP vaccinated group compared to the PBS+Alhydrogel® control group. This type of response has been observed in a previous study and was attributed to unique elevations in pro-inflammatory components, suggesting that cytokine storms may contribute to animal death (66). Renal changes were more seen in PBS+Alhydrogel® control group compared to the MCP vaccinated group. The mechanisms involved in the progression of severe disease in the MCP vaccinated group remain unknown and further studies are needed. We assume that higher levels of antibodies generated against MCP may be inducing antibody dependent enhancement (ADE) of infection and inflammatory response, which is well recognized in viral infections. Resistance to complement mediated killing is reported in *Leptospira*, enabling it to bypass this component of the innate immune system (83). Binding of antibodies can activate cytokine release and complement cascade for host protection, but when uncontrolled, can lead to tissue damage and inflammation (84). There has been some evidence of ADE associated with bacterial infections, including *Streptococcus*, *Acinetobacter*, *Neisseria* and *Pseudomonas*, and the proposed causes include virulence and adhesion enhancement, serum resistance, and protection from bactericidal killing (84). Some of the antibodies mentioned in the above report have been shown to inhibit the complement mediated killing of bactericidal sera, and this type of blocking antibodies are observed in some Gram-negative species, including those causing pyelonephritis (84). In our case, the anti-MCP antibodies may also block the MCP signaling cascade and may limit chemotaxis, preventing *Leptospira* from reaching its destination but facilitating its maintenance and replication in the blood stream. Further functional studies on MCP proteins might reveal some of these factors. We can speculate that in endemic areas, where humans are highly exposed to *Leptospira* antigens, the presence of blocking antibodies might lead to exacerbated clinical signs.

The histopathologic changes observed in the MCP vaccinated and control mice were different. MCP vaccinated group exhibited mild acute renal changes, in comparison to control group which had mild to moderate chronic changes including inflammation and fibrosis. Hepatic changes although subtle, were more pronounced in the animals with jaundice, which was present in more MCP vaccinated animals in comparison to controls. These observations support that in the MCP vaccinated animals, clinical signs are more severe and leads to a faster decline compared to the control group. In the PBS+Alhydrogel® control group more animals had chronic renal changes and slower health decline.

The animal challenges were performed by intraperitoneal route, a commonly used route in *Leptospira* challenge studies. This is not the natural route of *Leptospira* infection and can potentially bypass the initial innate immune mechanisms and overwhelm the host immune system and, hence, may underestimate the vaccine efficacy. The challenge dose, the age of the animal at challenge, and the challenge strain may also influence the outcome of vaccination challenge studies. Therefore, the standardization and development of uniform guidelines for challenge studies is also desirable.

## Conclusion

Vaccination with a recombinant methyl-accepting chemotaxis protein fragment did not protect mice from disease and renal colonization, despite eliciting a robust immune response. The vaccination appears to have enhanced the disease process in these animals. The role of this protein in *Leptospira* pathogenesis should be further evaluated to better understand the lack of protection or potential exacerbation of the disease process. The absence of immune correlates of protection from *Leptospira* infection is still a major limitation in the field and efforts to gather this knowledge is critically important.

## Author contributions

**Conceptualization:** Liana Nunes Barbosa, Sreekumari Rajeev

**Data curation:** Liana Nunes Barbosa, Sreekumari Rajeev

**Formal analysis:** Liana Nunes Barbosa, Alejandro Llanes, Kalvis Brangulis, Sarah C. Linn-Peirano, Sreekumari Rajeev

**Funding acquisition:** Sreekumari Rajeev

**Investigation:** Liana Nunes Barbosa, Sreekumari Rajeev

**Methodology:** Liana Nunes Barbosa, Kalvis Brangulis, Swetha Madesh, Bryanna Nicole Fayne, Sreekumari Rajeev

**Project administration:** Sreekumari Rajeev

**Resources:** Sreekumari Rajeev

**Supervision:** Liana Nunes Barbosa, Sreekumari Rajeev

**Validation:** Liana Nunes Barbosa, Sreekumari Rajeev

**Visualization:** Liana Nunes Barbosa, Sreekumari Rajeev

**Writing – original draft:** Liana Nunes Barbosa, Sreekumari Rajeev

**Writing – review & editing:** Liana Nunes Barbosa, Alejandro LIanes, Swetha Madesh, Bryanna Nicole Fayne, Kalvis Brangulis, Sarah C. Linn-Peirano, Sreekumari Rajeev

## Acknowledgments

We thank the University of Tennessee College of Veterinary Medicine, Center of Excellence in Livestock Diseases and Human Health for funding this project. We also thank Morris Animal Foundation (D21CA-080) for the funding to generate preliminary data. We are grateful to Mossman Laboratory Animal Facility staff and the student Assistant Ms. Cheri Bonnell for their support and assistance in this study.

## Supporting information

**S1 Table. Clinical score to assess the condition of mice infected with pathogenic *Leptospira*.** (.DOCX)

**S2 Table. Potential orthologs of MCP protein in 27 representative *Leptospira* genomes.** (.DOCX)

**S3 Table. Summary of results from individual mouse from the MCP vaccinated and PBS+Alhydrogel® control group.** (.DOCX)

